# The effects of crank power and cadence on muscle fascicle shortening velocities, muscle activations and joint-powers during cycling

**DOI:** 10.1101/2022.07.17.500375

**Authors:** Cristian Riveros-Matthey, Timothy J. Carroll, Glen A. Lichtwark, Mark J. Connick

**Author notes:** Corresponding author: Cristian Riveros-Matthey, School of Human Movement and Nutrition Sciences, University of Queensland, St Lucia Street, Queensland, 4072.

## Abstract

Whilst people typically chose to locomote in most economical fashion, during cycling on a bicycle they will, unusually, chose cadences that are higher than metabolically optimal. Empirical measurements of the intrinsic contractile properties of the vastus lateralis (VL) muscle during submaximal cycling suggest that the cadences that people prefer (i.e., self-selected cadences: SSC) allow for optimal muscle fascicle shortening velocity for the production of knee extensor muscle power. It remains unclear, however, whether this is consistent across different power outputs where SSC is known to might be affected. We examined the effect of cadence and external power requirements on muscle neuromechanics and joint powers during cycling. VL fascicle shortening velocities, muscle activations and joint-specific powers were measured during cycling between 60 and 120rpm (and the SSC), while participants produced 10%, 30%, and 50% of peak maximal power. VL shortening velocity increased as cadence increased but was similar across the different power outputs. Although no differences were found in the distribution of joint powers across cadence conditions, the absolute knee joint power increased with increasing crank power output. Muscle fascicle shortening velocities increase in VL at the SSC as pedal power demands increase from submaximal to maximal cycling. It therefore seems highly unlikely that preferred cadence is primarily driven by the desire to maintain “optimal” muscle fascicle shortening velocities. A secondary analysis of muscle activation patterns revealed that minimizing muscle activation is likely more important when choosing a cadence for given pedal power demand.

## Introduction

Humans tend to locomote in ways that minimize energy consumption (Bertram 2005; Ralston 1958; Hoyt 1981; Zarrugh, Todd, and Ralston 1974; Alexander McN. 1989; Alexander 2000; Minetti and Alexander 1997; Snaterse et al. 2011; Sparrow and Newell 1998). A good example of this is during walking or submaximal running (i.e., below lactate threshold), where people tend to locomote at a step frequency near that which minimizes the metabolic energy cost. However, people do not show the same behavior when riding a bicycle, and tend to use higher cadences than the metabolically optimal one (Brisswalter et al. 2000; Alejandro Lucia, Jesus Hoyos 2000; Anthony P. Marsh and Martin 1997; Brennan et al. 2019). Apparently, other criteria besides energetic economy are at play in the self-selected pedaling rate during cycling.

There are several alternative factors that have been proposed to influence cadence selection in cycling (Anthony P. Marsh, Martin, and Sanderson 2000; Brennan et al. 2019). For example, factors such as muscle activation (A. P. Marsh and Martin 1995; MacIntosh, Neptune, and Horton 2000; Neptune, Kautz, and Hull 1997; Sarre et al. 2003; Takaishi et al. 1996; Baum and Li 2003) and net joint moments (Neptune and Hull 1999; Hull and Gonzalez 1987; Anthony P. Marsh, Martin, and Sanderson 2000) were reportedly minimized at high cadences under submaximal conditions in experimental and simulated environments. However, most of these studies did not measure the SSC directly. For instance, by exploring the amount of muscle activation per pedaling cycle, studies have shown a minimization of EMG root mean square (EMG RMS) between 80 rpm (in vivo) (Takaishi et al. 1996) and 90rpm (in silico) (Neptune and Hull 1999). Importantly, the cadence at which EMG per cycle is minimized shifts towards higher cadences as power output increases (MacIntosh, Neptune, and Horton 2000). However, these protocols were tested under fixed cadence conditions, not accounting for a self-selected cadence protocol. Moreover, the study conducted by Marsh et al. (Anthony P. Marsh, Martin, and Sanderson 2000), which did include an SSC protocol, found only a modest association between the cadence selected by cyclists and the cadence that minimized a joint moment-based cost function. Thus, it remains unclear whether EMG and joint moment variables are the main drivers of the SSC.

A third mechanical factor that has recently drawn attention is related to the intrinsic contractile properties of the muscles. The ability of muscles to generate maximal force is critically influenced by the length and velocity of muscle fibers, factors that ultimately constrain the capacity of muscles to produce power during cycling. In this context, force-length and force-velocity properties and their mechanical function during cycling have been reported for the vastus lateralis (VL) muscle, a major power producing muscle, under sub-maximal conditions during cycling (Austin, Nilwik, and Herzog 2010; Muraoka et al. 2001; Brennan et al. 2018; 2019). For example, Muraoka et al., (Muraoka et al. 2001) found that the timing of shortening of fascicles varies compared to the muscle-tendon unit (MTU) because of interaction with the elastic tendinous tissue in series with fiber. Although, these observations were done at low cadences and power outputs, this suggests that the elasticity of the VL tendinous tissue may decouple fascicle shortening from that of the muscle-tendon unit. In the same line, Brennan and colleagues (Brennan et al. 2018), through ultrasound imaging, demonstrated that fibers operate close to their optimal length (*L_o_*) for maximal force-production and velocity (*V_o_*) to achieve maximum power output at different cadences (range from 40 to100rpm). They found that while VL fascicle shortening velocity increased with cadence, there was a plateau in average shortening velocity at high cadences. Additionally, they found that the operating velocities of the fascicles at the preferred cycling cadence was approximately equal to that predicted to maximize power according to the force-velocity relationship (Brennan et al. 2019). These findings provide a deeper understanding of how the muscle properties interact in a range of cadence conditions; however, they are limited by the fact that a single power output condition was assessed (2.5 W· *kg*^-1^). More data are required on the relationship between self-selected cadences and muscle fascicle shortening velocities across a range of external power requirements, where force requirements vary.

One potential strategy that the CNS can use, under varying cadences and power external requirements, is to change the distribution of the joint powers. For example, at submaximal conditions (80-340W), the knee extensors contribute the greatest proportion of total lower limb power, with smaller contributions from the knee flexor and hip extensor powers (Ericson 1988; Bini et al. 2010). Conversely, during cycling at near-maximal power outputs (350-850W), there is a shift in the joint power distribution of the lower limb, such that the hip extensor contribution to crank power is twice the knee extensor joint power contribution, and the knee flexor contribution becomes equal to the knee extensor contribution (Elmer et al. 2011; McDaniel et al. 2014). These responses may be influenced by the constraint of knee extensors to generate power in those conditions (e.g., due to high shortening velocities). Thus, triggering a redistribution of the power mainly driven by the hip. However, the question of how shortening velocities constrain muscle power capacity at high power requirements remains unexplored.

The aim of this study was to describe how muscle mechanics and joint-specific powers vary with crank power and pedaling cadence during cycling and to determine whether fascicle shortening velocity at the self-selected cadence changes with increasing power output. We also explored how muscle activation varies across crank power and cadence conditions. Based on the study by Brennan (Brennan et al. 2019), we hypothesized that there would be little change in the VL fascicle shortening velocity at the SSC for different external powers, which would support the idea that people prefer to cycle at cadences that maximize the power generating capacity of the muscles that contribute most strongly to the task. We expected that the power contribution of the knee extensors would also decrease with increased crank power requirements. To test these hypotheses, we used ultrasound to measure muscle fascicle length and velocity changes in the VL muscle, and instrumented pedals and motion capture to calculate joint powers by inverse dynamics. We tested the joint power distribution and work done by the lower limb to document the relative contributions of hip, knee and ankle joints across the different external power requirements. To study the interaction between the VL muscle fascicle behavior and adjacent structures at the knee joint, we also quantified electromyographic activity (EMG) from gluteus maximus (GMAX), vastus lateralis (VL), rectus femoris (RF), biceps femoris (BF) and semitendinosus (ST).

## Material and methods

### Subjects

Twelve healthy men from the university community and cycling sports clubs (mean and standard deviation, age= 27 ± 9.9 yr., mass= 70 ± 7.9 kg, height= 1.76 ± 0.054 m) participated in this study. We recruited level 3-4 cyclists, who are daily riders with a considerable history of involvement with the discipline. The participants delivered their informed written consent, and all procedures were approved by the Human Research Ethics Committees (HRECs) of the University of Queensland.

All cycling was performed on a bicycle ergometer (Excalibur Sport, Lode BV, Groningen, The Netherlands) that was adjusted according to the anthropometric characteristics of each subject. The saddle height was normalized to 100% of the greater trochanter length (Bini et al. 2014). The trunk angle was standardized according to the preference of each subject within the range of 35-45 °, with the hands placed on the handlebar drops. The angle of the trunk was defined by the line that connects the anatomical landmarks of acromion and greater trochanter with respect to horizontal. Visual feedback was used to ensure that posture was maintained during the experimental sessions. The crank length was set to 175 mm. Participants wore cleated shoes which clipped into the pedals (SH-R070, Shimano, Osaka, Japan; SH-R540, Shimano, Osaka, Japan).

### Experimental design

Each participant completed two assessment sessions on different days. The first day-assessment was divided into three different parts. The first part comprised the force-velocity test, in which the maximum power that the participant could exert on the ergometer cranks (Pmax) was measured (Fig. 1C). This test involved two trials, in each of which the participant performed a maximal sprint of 5s with a resistance of 1 *Nm*^*kg*-1^ body mass, followed by a rest period of 5 minutes (Sylvain Dorel et al. 2010; Morin and Samozino 2019; S. Dorel et al. 2005). Participants were verbally encouraged to perform maximally during each trial. Secondly, the self-selected cadence of each participant was determined at three different intensities (10%, 30% and 50% of the Pmax). These tests lasted for 5 minutes, 1 minute, and 30 seconds for the 10%, 30% and 50% intensities respectively (Fig. 1D). Finally, participants underwent a familiarization of the cadence-matching protocol to be applied on session-day two. Each participant completed 6 bouts of cycling at cadences that were randomly selected from 60 to 120rpm, and at power outputs that were randomly selected from 10%, 30% and 50% of the Pmax.

**Fig. 1.**
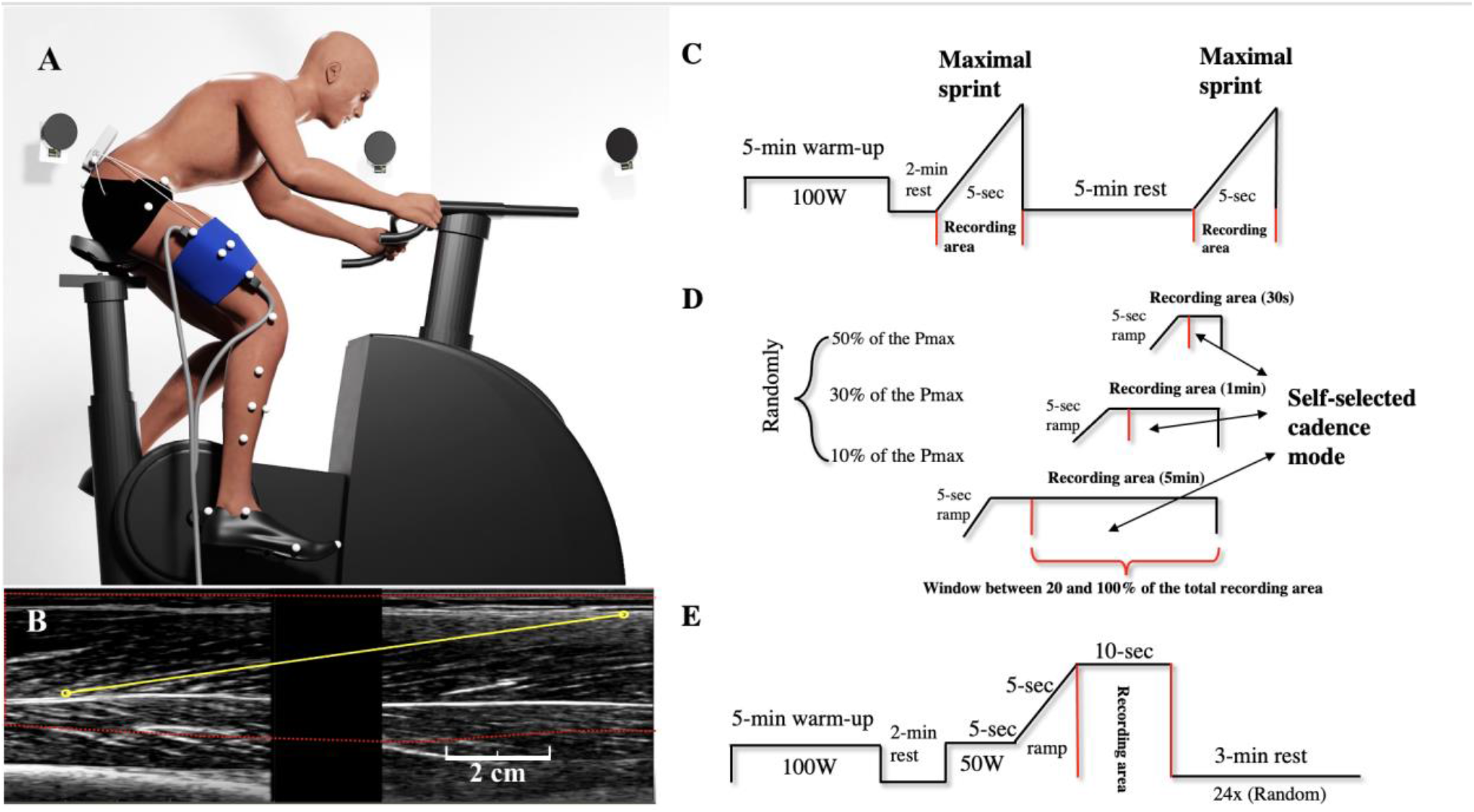
Experimental setup. Schematic representation of the experimental design, specifically associated with the second-day assessment, data collected from 3-dimensional motion-capture, ultrasound of the VL muscle, surface electromyography and radial tangential force on the cycle ergometer (A). Estimating fascicle length in VL muscle through dual probes images during cycling. (B). Illustrating the experiment protocol that is divided into two assessment days. The first day-assessment: force-velocity test (C), and self-selected cadence protocol (D). And the main cycling protocol conducted during the second day-assessment (E).

In the second-day assessment, each subject cycled at three different percentages of the maximum power obtained in the force velocity test described above. For each power level, participants were asked to cycle at eight different cadences (60rpm, 70rpm, 80rpm, 90rpm, 100rpm, 110rpm, 120rpm and the self-selected cadence), resulting in a total of 24 trials (Fig. 1E). Before the session, participants began with 5-m warm-up at 100 W at a self-selected cadence. For each trial, they were asked to cycle for approximately 20 seconds. For the first 5-s, riders pedaled at 100W at the nominated cadence. Then, in the remaining 15-s, participants were exposed to a 5-s incremental ramp in crank resistance until the desired power output was reached, at which intensity their behavior was recorded for 10-s. Each trial concluded with active recovery at 100W, for three, two or 1 minutes depending on the previous intensity (50%, 30%, 10% of the Pmax) (Wilkinson, Cresswell, and Lichtwark 2019). The following measurements were taken during each trial: 3-dimensional motion-capture, surface electromyography, ultrasound of the VL muscle, radial tangential force, and pedal angle on the cycle ergometer (Fig. 1A).

### Instruments

#### Kinematics

Lower limb kinematics were recorded at 200Hz using fourteen 3-dimensional motion-capture cameras (Oqus, Qualisys, AB, Sweden) and software (Qualisys Track Manager). Twenty-two small light-weight reflective markers were attached to participants’ skin. Markers were placed in rigid-clusters of three on the mid-thigh and mid-shank, and individually on the anterior and posterior iliac spines, greater trochanter, medial and lateral epicondyles, medial and lateral malleoli, calcaneus, first and fifth metatarsal heads. After placing the reflective markers on each cyclist, a trial was captured in a static standing position with their arms crossed. The static recording was then used for scaling purposes during data analysis. To create a global coordinate system, reflective markers in the ergometer cycle were placed on the back of the saddle and at the left and right rear support angle of the cycle ergometer. The cameras tracked the 3D trajectories from each marker at a 200Hz sampling rate.

#### External Forces

We used a wireless, instrumented crank device (Axis, SWIFT Performance, Brisbane, Australia) to measure the orthogonal crank forces in polar coordinates, crank torque (tangential force * length), and the crank axial force (radial force). From these measurements, the vertical (Fz) and horizontal (Fx) components of pedal forces were derived. Force data were synchronously sampled via an analog to digital (A/D) board capturing at 100 Hz via the Qualisys Track Manager Software. Before each assessment, dynamic and static calibrations were performed on the instrumented cranks.

#### Ultrasonography recording equipment

Two B-mode ultrasonography devices were used to assess VL muscle fascicle lengths (LV7.5/60/96Z, TELEMED, Vilnius, Lithuania) via two flat transducers mounted in-series within a frame. Before the recording, the transducers were covered in conductive gel and in direct contact with the participant’s skin. Then, both were placed in-series on the mid-thigh, taking as a reference a straight line between the greater trochanter and superior face of the patella (Brennan et al. 2018), and secured using a self-adhesive compression bandage (Fig. 1A). Both ultrasound units were synchronized utilizing analog pulses sent from a separate A/D board that triggered collection of each individual frame (Micro 1401-3; AnySyncro software). The sampling rate was configured and recorded, using Spike 2 software (Cambridge Electronic Design Ltd., Cambridge, England) through sending a 160 Hz quadratic logic pulse to the ultrasound equipment connected in parallel. This logic pulse also triggered the start of motion capture collection, allowing an exact synchronization between the probes and ultrasound recordings and the other signals.

#### Electromyography (EMG)

Surface EMG data (Myon 320 system; Myon AG, Baar, Switzerland) were collected at 2kHz using sensors placed on the GMAX, VL, RF, BF and ST of the right leg and were synchronously recorded with motion capture data via the A/D board controlled by the Qualisys Track Manager Software. The electrodes were adhered to the subjects’ skin following SENIAM guidelines with an interelectrode distance of 2cm (Hermens et al. 2000). Before the placement, the hair over the muscle was shaved with a disposable razor and the skin cleaned with 70% isopropyl alcohol.

### Data Analysis

#### Muscle fascicle shortening velocities

Images from the two transducers were concatenated considering a scale of 60 mm and with a space of 22 mm between each field of vision (FOV) of the probes, using a custom MATLAB script (Fig. 1B). A custom MATLAB script based on the Lucas-Kanade optical flow algorithm was used to track muscle fascicles and measure the changes in VL muscle fascicle length (Gillett, Barrett, and Lichtwark 2013; Brennan et al. 2018). The length changes were estimated considering a whole fascicle from the proximal intersection with the superficial aponeurosis and vice versa. The velocity of the VL fascicle respect to the crank angle was calculated as first derivative of their lengths with respect to time. The fascicle shortening velocity was calculated between the maximum and minimum length within the downstroke phase during crank angle (knee extension period). The peak and average shortening velocities during the downstroke phase were calculated.

#### Joint powers analysis

Data obtained from the forces recorded by the pedals of the instrumented crank and by the kinematic variables from 3-dimensional motion-capture cameras were exported and analyzed via a custom MATLAB script (R2019a, MathWorks Inc., USA). All signals were filtered using a zero-lag low pass second-order Butterworth filter with a 12 Hz cut-off frequency. To represent the task cycle, the crank angle was reconstructed from the right foot kinematic data from a virtual marker representing the pedal axle, considering markers from lateral, medial ankle and calcaneus. Also, this virtual marker was used to convert the tangential and radial forces into vertical and horizontal components with respect to the global coordinate system. To rotate the force components to the ergometer’s reference system, matrix rotation algorithms were applied, obtaining the resulting pedal forces, which were projected to the pedal axle. Subsequently, an open-source refined model of the lower extremity (Lai, Arnold, and Wakeling 2017) was scaled to each cyclist’s anthropometric measurements. Then, the cyclist-specific models, as well as the experimental kinematic and kinetic data were used to compute joint angles and joint moments applying inverse kinematic and inverse dynamic tools in OpenSim. We further calculated joint powers as the product of net joint moments and joint angular velocity. Joint work per cycle was calculated as the time integral of joint power per cycle. To calculate the joint-specific powers, flexion and extension phases were determined from the angular velocity signs. As such, negative specific powers were obtained when the moments and angular velocities were in opposite directions. Joint-specific powers were averaged over each crank cycle (Elmer et al. 2011).

#### Complementary variable analysis

All EMG signals were processed with a 15/500 Hz bandpass filter to remove non-physiological signal noise. To remove the analog channels offset, the signal median activation for each muscle was subtracted. The root mean square (RMS) was calculated with a moving window width of 50ms as an index of the signal amplitude. Further, the mean EMG signals for each muscle were normalized to the peak EMG RMS value in each cycle during the 90rpm cadence at 50% Pmax. Some EMG data of VL muscle was discarded due to sensor displacement elicited by contact with ultrasound transducers. EMG data collected from GMAX and RF muscles were averaged across 12 subjects, whereas VL, BF and ST were averaged across 10 subjects.

To have a better representation of the changes in the joint moments, the averaged value of the knee joint moment in the positive extension phase was considered. The positive extension phase was calculated by determining period of positive knee angular velocity for each cadence condition. The lengths and velocities of the VL MTU were calculated using the inverse kinematics results and the muscle analysis tool in OpenSim.

### Statistical analysis

All values were taken as the average of the first five cycles of the right crank per each cadence condition and crank power requirement (n.b. the crank angle at top-dead-center was defined as 0deg). A two-way repeated measures ANOVA was used to test for main effects of cadences and crank power requirements and interaction effects on muscle shortening velocities, joint powers, positive knee extensor moments and EMG signals. A repeated measures one-way ANOVA was used to test the main effect of the crank power requirement at SSC on muscle shortening velocities.

Tukey multiple comparisons tests were made across cadences conditions within each level of analysis. To quantify how much more likely the muscle shortening velocities at SSC across crank power requirements are under the null hypothesis, we performed Bayesian ANOVA analyses, considering a credible interval of 95% and post hoc tests. The Bayes factor towards the null hypothesis was denoted as *BF*_01_. For its interpretation, *BF*_01_ >1 and <3 there is *anecdotal* evidence, *BF*_01_>3 and <10 *moderate* evidence, *BF*_01_>10 and <30 *strong* evidence, *BF*_01_>30 and < 100 *very strong* and *BF*_01_>100 extreme towards the null hypothesis. The EMG data were also fitted through a second-order polynomial nonlinear regression to examine the relationship between the mean EMG RMS activation and cadence at each pedal power requirement. Coefficient of determination (*R*^2^) was used to compare the experimental and predicted SSC in which the minima EMG RMS was elicited across muscles and pedal power requirements. An α level of 0.05 was set for all tests. All data were processed using Prism 7 (GraphPad Software Inc., La Jolla, CA) and JASP (JASP Team, 2019).

## Results

During the force-velocity test, participants performed a total averaged power of 898 ± 37 W. This yielded power targets at 10% (89.8 ± 3 W), 30% (270 ± 10 W) and 50% (449 ± 18 W) of Pmax. The self-selected cadence protocol showed that cyclists increased their cadences as crank power increased (e.g., 30% versus 50% of the Pmax *P* = 0.001), reaching cadences of 94 ± 11, 103 ± 11 and 117 ± 12 rpm at 10%, 30% and 50% of the Pmax respectively.

### Muscle fascicle shortening velocities

There was no main effect of pedal power requirements on mean fascicle shortening velocities (*F*_(1,15)_= 0.23, *P* = 0.71, *η*^2^ = 0.007). There was a significant effect of cadence on mean fascicle shortening velocity, with mean fascicle shortening velocity increasing with cadence (*F*_(2,24)_= 111, *P* = 0.001, *η*^2^ = 0.60) in a similar manner across all crank power requirements (Fig. 2). The increase in mean fascicle shortening velocity with cadence was approximately 1.01, 0.92 and 1.16 cm/s per 10rpm at 10%. 30% and 50% of Pmax power conditions respectively. There was no interaction effect between pedal power requirements and cadence on mean fascicle shortening velocity (*F*_(4,52)_= 0.62, *P* = 0.66, *η*^2^ = 0.04).

**Fig. 2.**
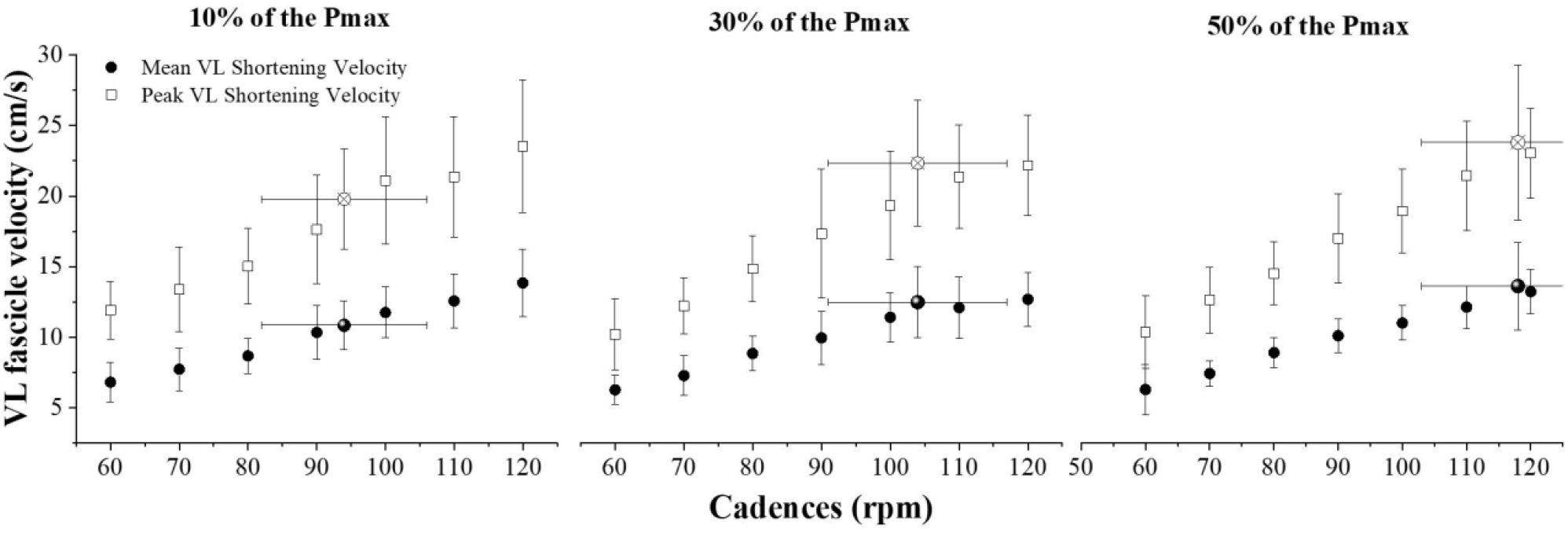
Mean (black dot) ± SD and Peak (empty square) ± SD of VL fascicle shortening velocities absolute values at 60, 70, 80, 90, 100, 110, 120rpm and SSC at different external power requirements. The self-selected cadence (SSC) is 94rpm at 10% of the Pmax, 103rpm at 30% of the Pmax, 117rpm at 50% of the Pmax (for means: a highlighted black dot; for peaks: empty squares).

There was a significant main effect of pedal power requirement on mean fascicle shortening velocity at SSC (*F*_(2,33)_= 3.2, *P* = 0.04, *η*^2^ = 0.83) 10% of Pmax = 10.8 ± 1.8 cm/s; 30% of Pmax = 12.5 ± 2.5 cm/s; 50% Pmax = 13.5 ± 3.1 cm/s). Post hoc comparison showed a significant increase in fascicle velocity between 10% and 50% Pmax conditions (*P* = 0.04), but not between 10% and 30% of Pmax (*P* = 0.26) and 30% and 50% of Pmax conditions (*P* = 0.61) (Fig. 2). The Bayesian ANOVA post hoc analysis showed *anecdotal* evidence inclined to *H*_0_ when evaluating mean shortening velocities at SSC (e.g., 10% versus 30% of Pmax: *BF_01_*= 0.8; 30% versus 50% of Pmax: *BF*_01_=2).

There was also no significant effect of pedal power requirement on peak fascicle shortening velocity (*F*_(1,14)_= 0.34, *P* = 0.62, *η*^2^ = 0.13; Fig. 2). There was a significant effect of cadence on peak fascicle shortening velocity (*F*_(2,27)_= 94, *P* = 0.001, *η*^2^ = 0.58), with shortening velocity increasing with increasing cadence (Fig. 2). The was no significant interaction effect between crank power requirement and cadence on peak fascicle shortening velocities (*F*_(4,48)_= 0.55, *P* = 0.72, *η*^2^ = 0.04).

There was no significant main effect of pedal power requirement on peak fascicle shortening velocity at SSC (*F*_(2,33)_= 2.2, *P* = 0.12). *Modest* evidence in favor of *H*_0_ was found on peak shortening velocities at SSC across pedal power requirements (e.g., 10% versus 30% of Pmax: *BF_01_*= 1.2; 10% versus 50% of Pmax: *BF*_01_= 0.5; and 30% versus 50% of Pmax: *BF*_01_= 2).

### Joint powers and mechanical work

The power output for each joint relative to the total summed power across joints (for all crank power requirements and cadence conditions) is shown in Fig. 3A, whilst the individual contributions of positive and negative powers for flexion and extension phases are shown for the hip (Fig. 4A), knee (Fig. 4B) and ankle (Fig. 4C). Fig. 3B illustrates the total summed mechanical work of each joint across all cadences and crank power requirement conditions.

**Fig. 3.**
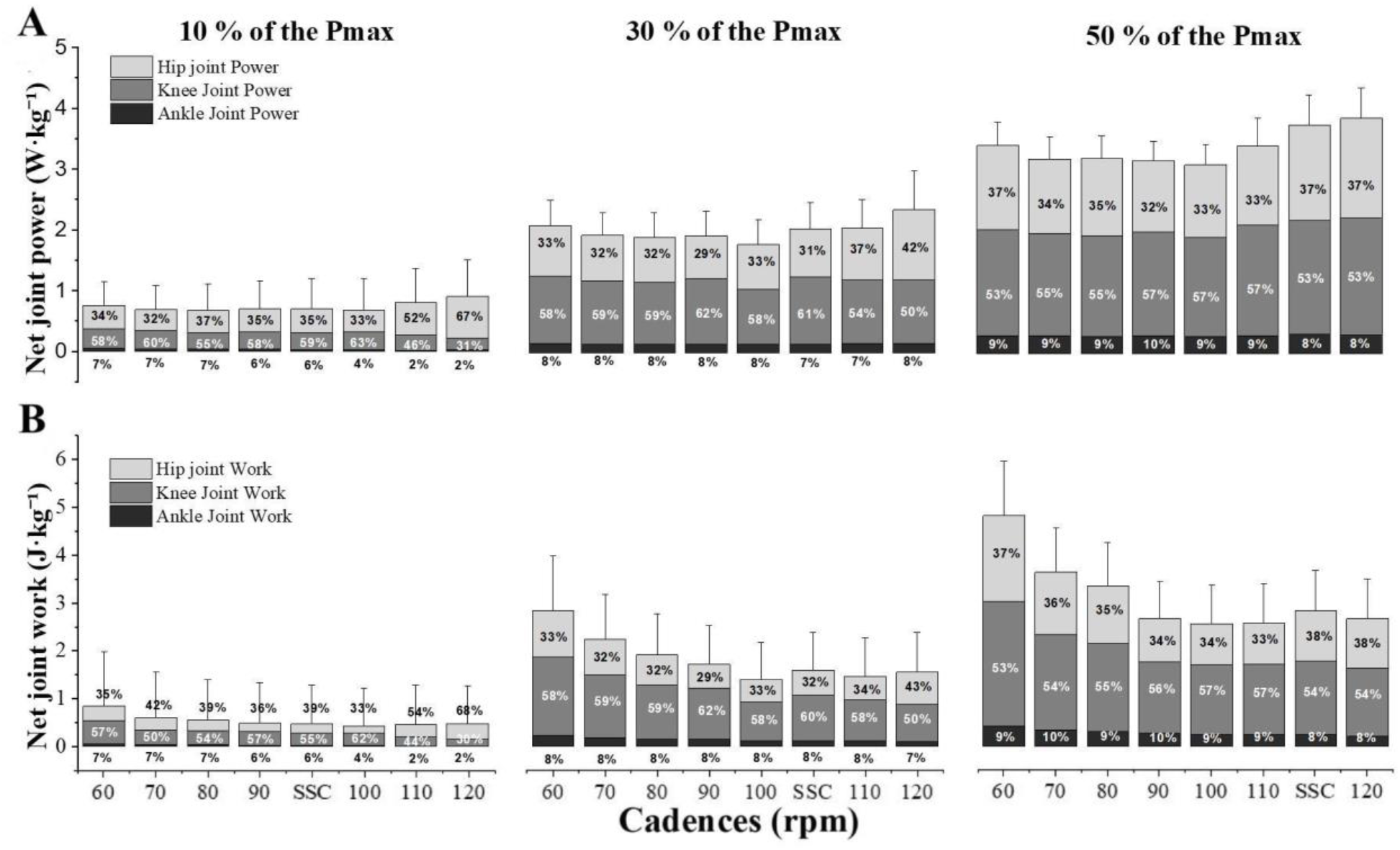
Joint power distributions (means ± SD) (A) and joint work distribution (means ± SD) (B) at 60, 70, 80, 90, 100, 110, 120rpm and SSC at different external requirements. The SSC is 95rpm at 10% of the Pmax, 103rpm at 30% of the Pmax, 113rpm at 50% of the Pmax.

**Fig. 4.**
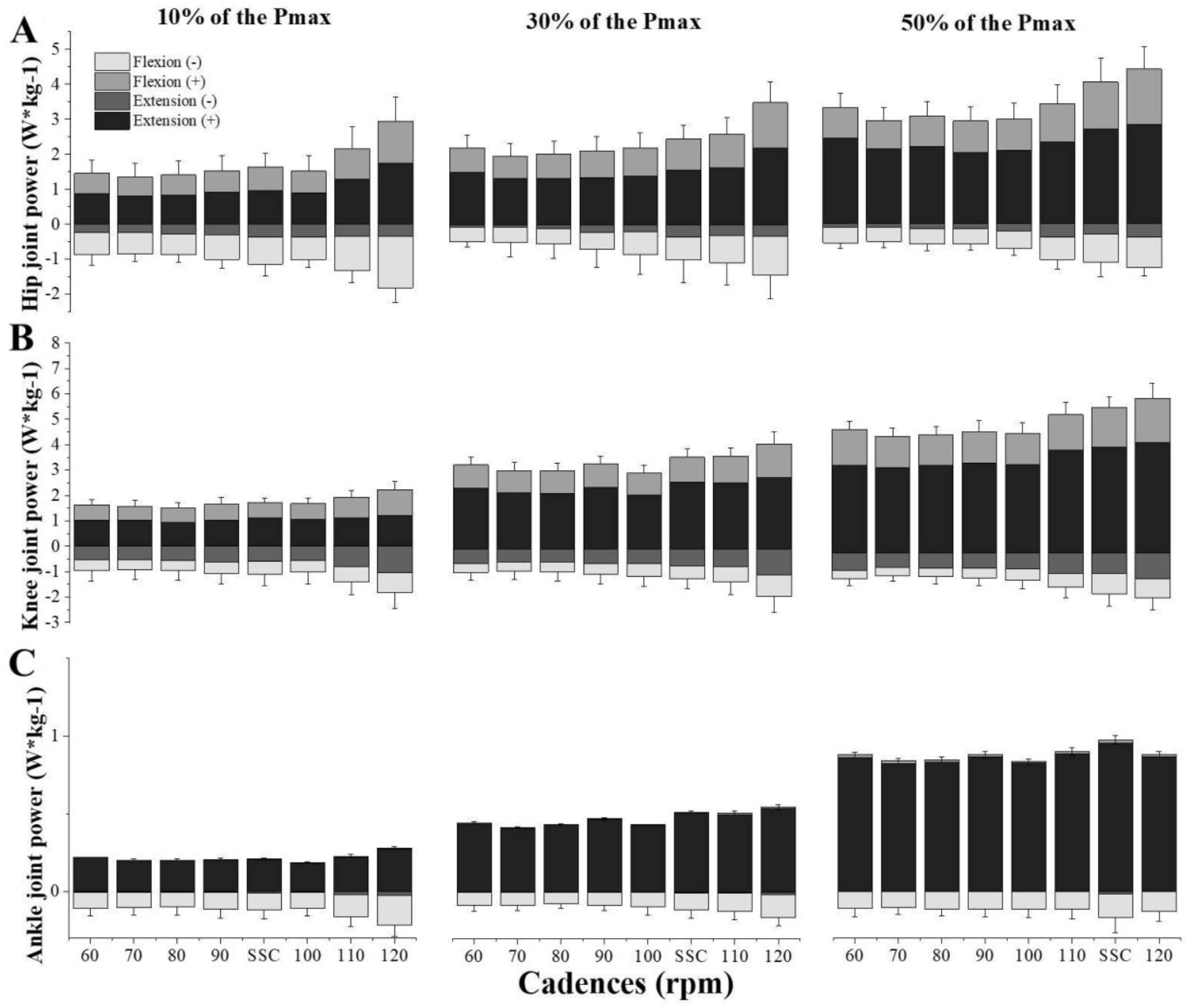
Decomposition of joint power (A to C) into positive and negative contributions (means ± SD) during net flexor and extensor muscle powers of hip, knee and ankle at 60, 70, 80, 90, 100, 110, 120rpm and SSC at different external requirements. The SSC is 95rpm at 10% of the Pmax, 103rpm at 30% of the Pmax, 113rpm at 50% of the Pmax.

A main effect of crank power requirement (hip joint: *F*_(1,14)_= 22, *P* = 0.002, η^2^ = 0.24; knee joint: *F*_(1,13)_= 128, *P* < 0.001, η^2^ = 0.60), cadence (hip joint: *F*_(2,28)_= 13, *P* < 0.001, η^2^ = 0.04; knee joint: *F*_(2,32)_= 4, *P* = 0.01, η^2^ = 0.06) and an interaction effect (crank power requirements x cadence) (hip joint: *F*_(4,45)_= 5, *P* = 0.009, η^2^ = 0.09; knee joint: *F*_(3,35)_= 3, *P* = 0.049, η^2^ = 0.04) were found for the relative contribution of hip and knee joint power to total power. Ankle joint power was affected by crank power requirement (*F*_(1,11)_= 49, *P* < 0.001, η^2^ = 0.42), but there was no cadence (*F*_(3,40)_= 1.4, *P* = 0.24, η^2^ = 0.05) or interaction effects (*F*_(2,23)_= 0.35, *P* = 0.72, η^2^ = 0.01). All joints increased their power contribution with increased crank power requirement (Fig. 3A). A main effect of cadence shows that there was a greater summed power output across hip and knee joints powers at higher cadences. However, this was mainly constrained to cadences above 110 rpm (e.g., hip joint power at 90rpm (0.37 W·*kg*^-1^; 0.70 W·*kg*^-1^; 1.17 W·*kg*^-1^) vs 110rpm (0.53 W·*kg*^-1^; 0.85 W·*kg*^-1^; 1.30 W·*kg*^-1^): *P* = 0.01, *P* = 0.02 and *P* = 0.01 at 10%, 30% and 50% of the Pmax respectively; Fig. 3A). Breaking down power into flexion and extension components saw similar trends: hip extension (*F*_(3,33)_= 34, *P* < 0.001, η^2^ = 0.09), knee extension (*F*_(2,26)_= 50, *P* < 0.001, η^2^ = 0.02) and knee flexion powers (*F*_(2,21)_= 29, *P* < 0.001, η^2^ = 0.13) (Fig. 4B) increased in unison as power increased.

A significant main effect of crank power requirement (hip joint: *F*_(1,18)_=132, *P* < 0.001, η^2^ = 0.15; knee joint: *F*_(1,15)_= 311, *P* < 0.001, η^2^ = 0.51; ankle joint: *F*_(1,12)_= 85, *P* < 0.001, η^2^ = 0.056), cadence (hip joint: *F*_(1,18)_= 17, *P* < 0.001, η^2^ = 0.02; knee joint: *F*_(2,28)_= 26, *P* < 0.001, η^2^ = 0.08; ankle joint: *F*_(2,30)_= 34, *P* < 0.001, η^2^ = 0.08) and interaction effect (hip joint: *F*_(5,57)_= 17, *P* < 0.001, η^2^ = 0.001; knee joint: *F*_(2,27)_= 7, *P* < 0.001, η^2^ = 0.002; ankle joint: *F*_(4,46)_= 9, *P* < 0.001, η^2^ = 0.002) was found for hip, knee, and ankle joint work. A significantly larger work per cycle was found at low cadences across hip and knee joints (e.g., knee joint work at 60rpm (0.48 J·*kg*^-1^; 1.64 J·*kg*^-1^; 2.50 J·*kg*^-1^) vs 80rpm (0.29 J·*kg*^-1^; 1.13 J·*kg*^-1^; 1.84 J·*kg*^-1^); *P* = 0.004, *P* = 0.001 and *P* = 0.001 at 10%, 30% and 50% of the Pmax respectively; Fig. 3B). Nonetheless, the distribution of work across the three joints remained similar across cadences for each of the power conditions tested.

### Complementary variables

There was a main effect of crank power requirement (*F*_(1,15)_= 269, *P* < 0.001, η^2^ = 0.48), cadence (*F*_(3,38)_= 16, *P* < 0.001, η^2^ = 0.01) and their interaction (*F*_(5,58)_= 2, *P* = 0.02, η^2^ = 0.004) on positive knee extensor moments, showing a systematic decrease as the cadence increased across all crank power output conditions (e.g., at 30% of the Pmax, 25.5 ± 10Nm, 23.9 ± 10Nm and 23.2 ± 10Nm for 60rpm, 70rpm and 80rpm versus 18.4 ± 10Nm at 120rpm; *P* = 0.001; *P* = 0.004; *P* = 0.001; Fig. 5B).

**Fig. 5.**
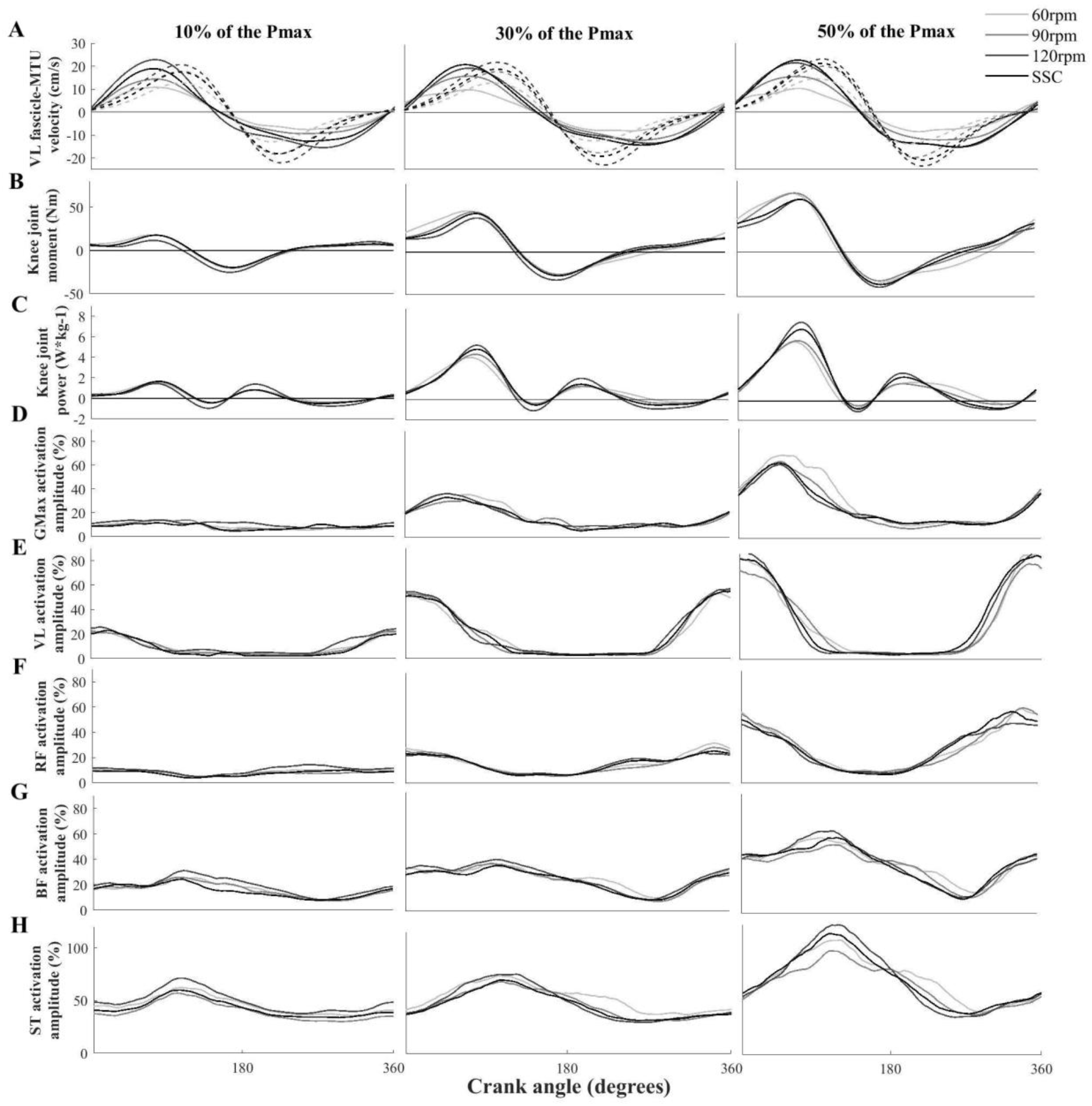
Mean waveforms of knee muscle mechanics and activation and gluteus maximus. Mean VL fascicle and MTU shortening velocities (A). Knee joint moments (B) and powers (C). Muscle EMG activation from GMAX (D), VL (E), RF (F), BF (G) and ST (H). All the waveforms were plotted across 60, 90, 120rpm and SSC at 10%, 30% and 50% of the Pmax against the crank angle cycle. The crank angle is 0 at top-dead-center. The SD and remaining cadences (70, 80, 100, 110rpm) were omitted for clarity.

A significant main effect of crank power requirement (GMAX:*F*_(1,14)_= 192, *P* < 0.001, η^2^ = 0.50; VL: *F*_(1,11)_= 777, *P* < 0.001, η^2^ =0.82; BF: *F*_(1,10)_= 210, *P* < 0.001, η^2^ = 0.68; ST: *F*_(1,14)_= 71, *P* < 0.001, η^2^ = 0.55), cadence (GMAX:*F*_(3,38)_= 8, *P* < 0.001, η^2^ = 0.01; VL: *F*_(1,16)_= 6, *P* = 0.008, η^2^ = 0.01; BF: *F*_(2,23)_= 3, *P* < 0.001, η^2^ = 0.09 and ST: *F*_(2,25)_= 3, *P* = 0.001, η^2^ = 0.02), and an interaction effect (GMAX: *F*_(3,38)_= 4, *P* = 0.007, η^2^ = 0.01; VL: *F*_(3,34)_= 3, *P* = 0.001, η^2^ = 0.05; BF: *F*_(3,31)_= 4, *P* = 0.01, η^2^ = 0.001; ST: *F*_(2,25)_= 3, *P* = 0.04, η^2^ = 0.001) was found for the mean EMG RMS for the GMAX, VL, BF and ST muscles. RF only was affected by crank power requirements (*F*_(1,17)_= 255, *P* < 0.001, η^2^ = 0.74) and cadence (*F*_(2,30)_= 3, *P* = 0.002, η^2^ = 0.06). The mean EMG RMS signal of all muscles increased as power pedal requirements increased (*P* = 0.001). Changes in muscle activation with cadence for different crank power requirements were non-linear for all muscles analyzed (Fig. 6). The nonlinear regression revealed a ‘U’ relationship between the mean EMG RMS and cadence for all muscles, showing a minimum at higher cadences as the power demands increased (Fig. 6). Furthermore, the mean EMG RMS activations at the SSC showed closeness to those where the local minima were predicted from the nonlinear regression for VL (F^2^ = 0.58 at 10% of the Pmax; *R*^2^ = 0.59 at 30% of the Pmax), BF (F^2^ = 0.67 at 10% of the Pmax only) and ST (F^2^ = 0.65 at 10% of the Pmax; *R*^2^ = 0.68 at 30% of the Pmax) muscles at 10% and 30% of Pmax (Fig. 7).

**Fig. 6.**
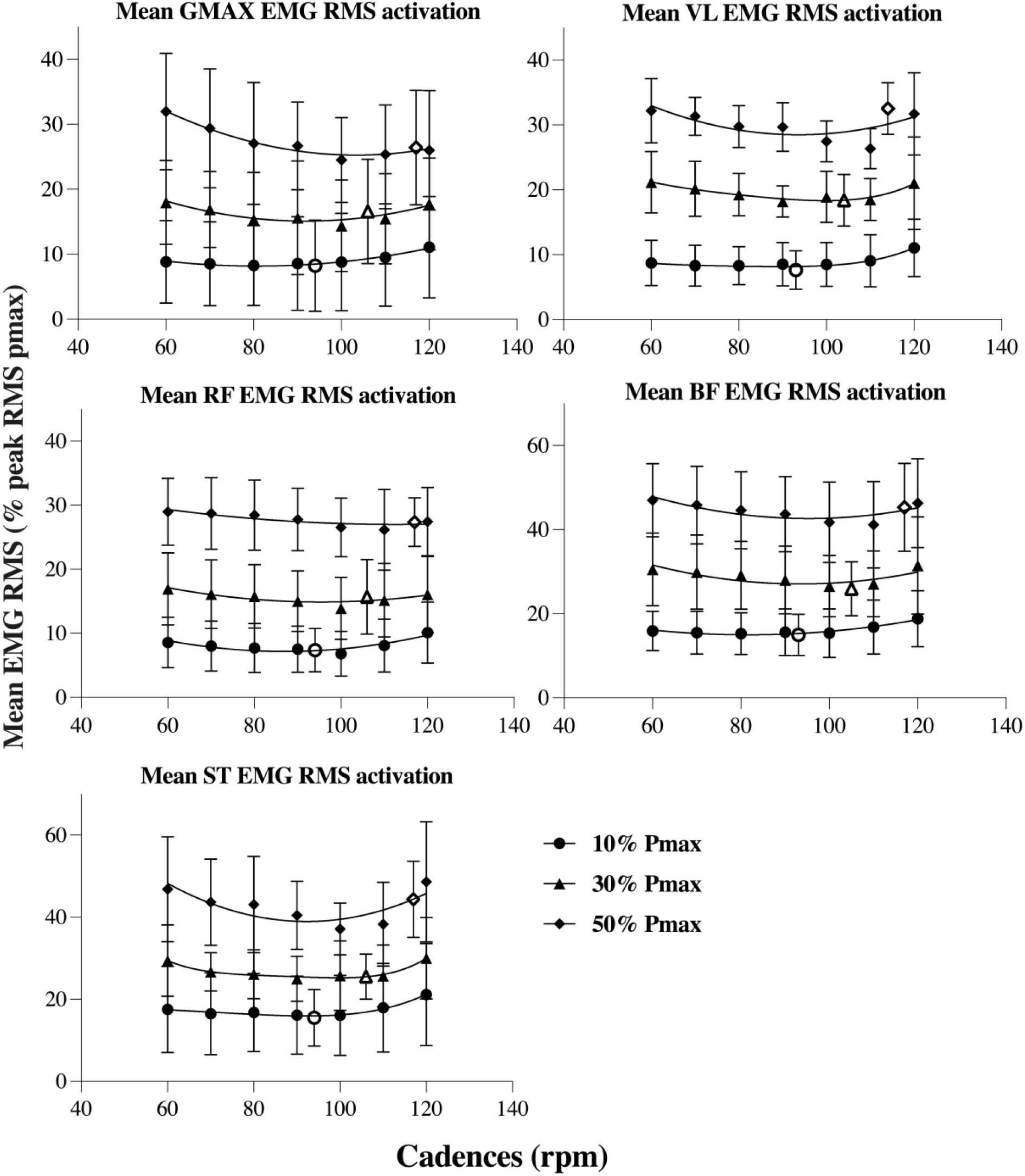
Mean EMG RMS of GMAX, VL, RF, BF and ST muscles at 60, 70, 80, 90, 100, 110, 120rpm and SSC at 10% of the Pmax (a highlighted black dot), 30% of the Pmax (a highlighted black triangle) and 50% of the Pmax (a highlighted black diamond). Data shown are means ± SD. Lines represent a second-order polynomial nonlinear regression. The SSC is 94rpm at 10% of the Pmax (empty circle), 106rpm at 30% of the Pmax (empty triangle), 117rpm at 50% of the Pmax (empty diamond).

**Fig. 7.**
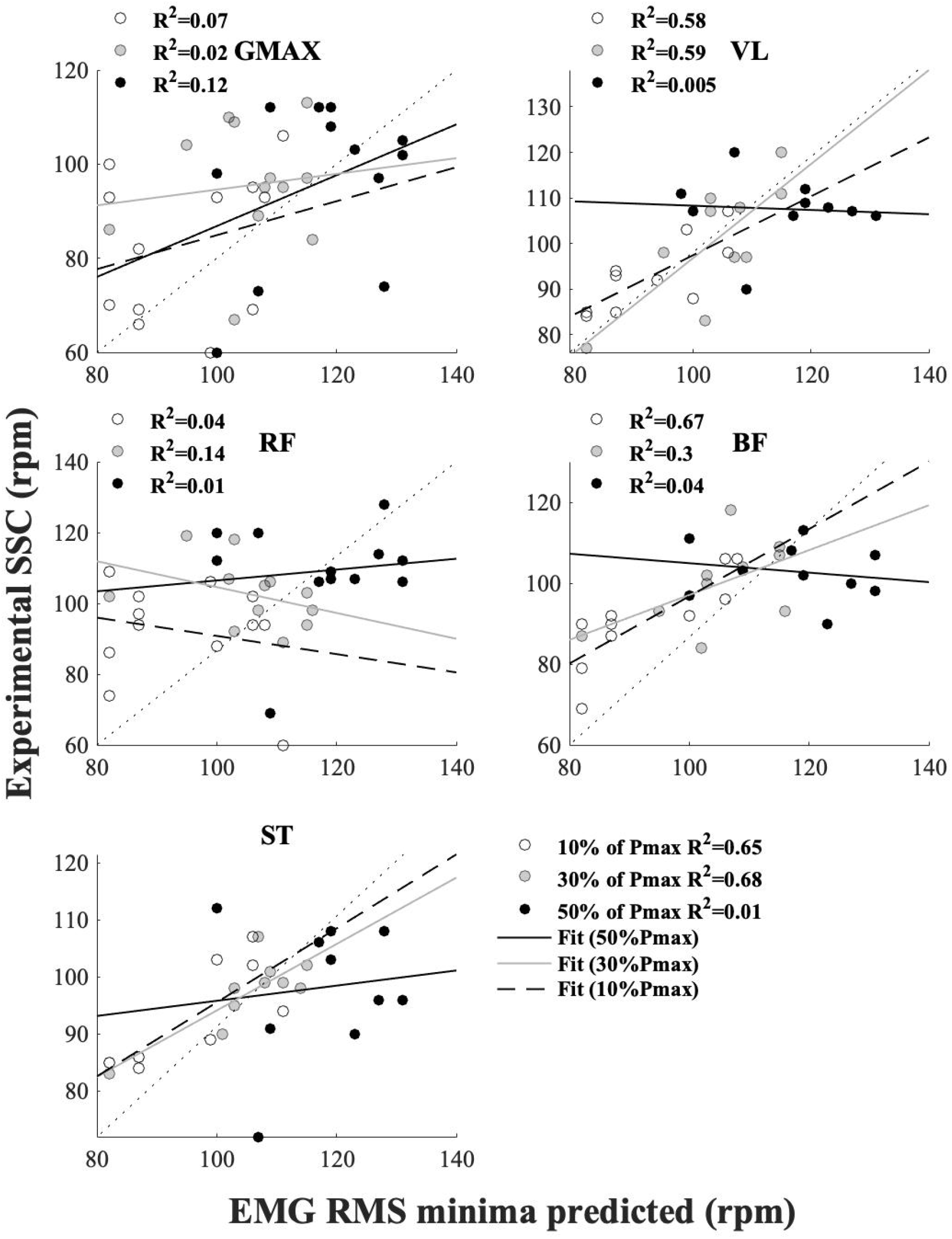
Linear regression and *R*^2^ coefficients between experimental SSC (rpm) and cadences where EMG RMS minima predicted were observed from the nonlinear regression per subject across muscles at 10% (empty dots), 30% (grey dots) and 50% (black dots) of the Pmax. Fit per pedal power demands are represented by discontinuous lines (10% of the Pmax), grey lines (30% of the Pmax) and black lines (50% of the Pmax).

## Discussion

This study examined changes in VL fascicle shortening velocity and joint-specific power contributions during cycling at a range of power outputs and cadences. We found that the net fascicle shortening velocities at a fixed range of cadences are not affected by changing the crank power requirement – hence fascicle velocity requirements largely mirror those of the MTU, which are constrained by the prescribed cadence. Our results also showed an increase in mean muscle shortening velocity at SSC as power requirement increased. We therefore reject our main hypothesis that self-selected cadence would have a similar VL fascicle shortening velocity, which has been shown to maximize power output (Brennan et al. 2019). It therefore seems highly unlikely that the cadence selection is driven by the desire to maintain an “optimal” shortening velocity. Our data also demonstrate that the power and work output of each joint increased as required; but showed little effect on their redistribution from one joint to another as pedal power requirements increased. This indicates that there was little change in motor coordination strategy across the range of power requirements studied here. Finally, we also found that the mean EMG RMS activations of knee muscle at the SSC are close to a local minimum at low pedal power demands, providing further evidence that the drive required to the muscle may be a key factor in SSC selection. Therefore, we suggest that it is unlikely that a mechanical performance criterion based on fascicle shortening velocities is an influential part of a cost function for bicycling; instead, muscle activations seem more promising when considering optimality factors that drive the SSC.

While fascicle shortening velocities in pre-set cadence conditions were not affected by changes in pedal power requirements, during the cadence selected by riders, our results suggest that VL fascicle shortening velocity increases as crank power requirements increase. Because SSC increased to higher cadences as power demands increased, this required an increase in VL fascicle shortening velocity, as reflected in our fixed cadence data. For example, SSC increased from 94 ± 11 rpm to 117 ± 12 rpm, in unison with the mean fascicle shortening velocity, which was 10.85 to 13.5 cm/s between 10% and 50% of Pmax, respectively (Fig. 2). Thus, it appears that there is no unique muscle shortening velocity that underpins a desire to maintain it optimal for maximal power generation. By contrast, previous findings suggested selecting high cadences at high power outputs may reduce the required force, generating a negligible impact on the net fascicle shortening velocity because of the series elastic structure involvement; maintaining the shortening velocities constant for power production capacity (Brennan et al. 2019). Our results do not support this assertion. Therefore, the increase in shortening velocity with cadence, irrespective of power requirement, provides evidence that the motor coordination strategy that governs during cycling may be driven by factors other than fascicle dynamics. However, the dynamics of the fascicle, along with the force requirements at each cadence, will subsequently influence the required activation of muscles, and therefore should be considered when exploring how the nervous system selects a preferred cadence.

The increased VL fascicle shortening velocity with cadence reduces the force generating potential of the muscle, however increasing cadence also reduced the knee extensor force requirements in all power conditions, and these two factors likely trade-off to influence the required VL muscle activation. Our VL EMG results show different interactions between cadence and activation under each power condition. As such, the u-shaped relationship between cadence and activation across power output conditions (Fig. 6), had a minimum that occurred at progressively increasing cadence, in line with the shift in SSC. Given that the mean VL fascicle shortening velocity for a given cadence was consistent across power conditions, the reduced activation at higher cadence and power combinations is likely related to subtle changes in the timing and magnitude of force production and subsequent power production. However, it is difficult to quantify individual muscle forces using inverse dynamics alone, and further simulation is likely required to better understand the relationship between activation and mechanical output.

Contrary to our expectation, we found that there was little change in power distribution across joints with changes in pedal power demands across the cadence range. This result contrasts with that of Elmer (Elmer et al. 2011), who found a redistribution at higher power outputs; specifically, a drop in knee extensor power (4%) and increase in hip power (4%). Our results suggest that at 90rpm (same as Elmer et. al.) changes in power requirement between 30% (270 ± 10W) and 50% (450 ± 18W) of Pmax, hip and knee extensor powers increased by 33% (from 1.3 to 2W; Fig. 4A) and 30% (between 2.4 and 3.5W; Fig. 4B), and the knee joint contributing approximately ~62-57% of the total joint power in those conditions (Fig. 3A). It appears that knee extensor power contribution is not affected by pedal power demands, increasing in magnitude as required from submaximal to maximal cycling. Although the knee extension force (as knee extension moment) decreased as the cadence increased (Fig. 5B), it increased from 23 to 34Nm between 30% and 50% of the Pmax at 90rpm. Thus, this suggests that it is unlikely that knee extensor power may be limited by force generation capacity and, hence, by the fascicle shortening velocity at high power demands.

Our results suggest that there may also be some further negative mechanical effects in selecting higher cadences as pedal power requirements increase. We found that at high cadences (> 100 rpm), the faster limb movements required generation of additional segmental kinetic energy, contributing to increases in total power production (Figs. 3A, 5C). This is likely due to the internal work generated to control higher segmental accelerations at faster cadences (Kautz and Neptune 2002). However, high cadences also affect the cyclist’s ability to effectively accelerate the crank due to the duration time reduction of the active muscles (Fig. 5 D, E, F, G, H; Fig. 6), generating an increase of knee negative power, specifically at the end of the upstroke phase in the crank cycle (Fig. 5C) (Neptune and Herzog 1999). In order to generate required knee flexor forces at the top of the crank cycle, our VL EMG data shows it is necessary to activate the VL muscle earlier relative to the crank position (Fig. 5E), but a similar time before this position. As such, the knee extensor moments increased earlier in the upstroke as cadence increased (Fig. 5B), contributing to greater absorption of energy at the knee.

In line with previous research (MacIntosh, Neptune, and Horton 2000), our study showed that muscles other than the VL (GMAX, BF, ST and RF EMG RMS) also exhibited a U’ shape relationship with cadence, with the minimum shifting to higher cadences as the crank power requirement increased (Fig. 6). These results suggest that it might be convenient to select higher cadences as the power demands on the pedal increase in order to maintain minimum muscle activation. As such, analyzing the EMG RMS at SSC, VL, BF and ST muscles showed a minimization of the EMG RMS close to the SSC whereas at 10% and 30% of the Pmax (Figs. 6, 7). By contrast, although the cadence selected by cyclists was larger at 50% than 10% and 30% of the Pmax, it exceeds the cadence with the minimum EMG RMS activation in all the muscles assessed (Figs. 6, 7). This finding might suggest that the cadence selected by riders at submaximal power output (10% and 30% of Pmax) may be a strategy to minimize fatigue, as previous studies suggest in simulated (Neptune and Hull 1999) and endurance cycling (Takaishi et al. 1996). These responses may be seen as the nervous system prioritizing the minimization of activation of overburdened muscles to prolong the movement duration (McDonald et al. 2022). While at high power pedaling demand landscapes, it appears that other factors might be involved besides minimizing muscle activation. For example, at very high-power outputs (near capacity), very high cadences may be preferable to minimize the amount of crank torque required. Alternatively, because our study solely contemplated muscles around the knee, it may be suggested that the SSC during submaximal cycling (10% and 30% of Pmax) may favor the resultant force on the pedal (Raasch and Zajac 1999; Patterson and Moreno 1990). This is based on the functions of monoarticular (VL) and biarticular (BF and ST) muscles in which BF might contribute to counteracting the forward component generated by VL muscle on the pedal, especially on the second part of the downstroke phase. Thus, the SSC may contribute to reducing the ineffective pedal forces, which are commonly shown at the bottom of the crank angle, expressing a reduction of muscle coactivation, and hence EMG RMS minimization. Therefore, it may be important to expand the analysis to a larger group of muscles in order to elucidate whether the sum of EMG RMS might be a local or global driver to consider when selecting a cadence.

### Limitations

Certain limitations should be acknowledged in our study. Only men were recruited to reduce the outcome variability. It has been shown that male muscles have higher power outputs and faster muscle groups (Glenmark et al. 2004), and hence likely different muscle fascicle shortening velocities. The SSC used as a preset value on the second day of assessment was based on the results obtained during the first day. Although SSC might be affected on the second day, the available evidence indicates that SSC is minimally affected by internal and external conditions during cycling (Hansen and Ohnstad 2008). Further, conditions such as ergometer configuration and power requirements were maintained on both days.

Our measurements of VL fascicle length and velocity based on ultrasound images have some potential limitations. For example, muscle deformation and horizontal shifting of connective tissues can cause an underestimation error of the fascicle length given by the muscle deformation (Brennan et al. 2017) (e.g., the fascicle origin and insertion), a greater horizontal shifting was observed on connective tissues in parallel (e.g., superficial, and deep aponeurosis) at lower crank power requirements (10% of the Pmax), which may affect the fascicle endpoints estimation. To counter this, aside from including ultrasound dual probes, the VL fascicle tracking was optimized by increasing the sampling rate to 160 Hz. This is because higher sample rates perform better for faster movements such as cycling due to the maximization in time resolution, enhancing the fascicle tracking (Cronin and Lichtwark 2013). Thus, the sampling rate selected likely improved the measurement of data extracted from the muscle fascicle.

## Conclusion

We found that while net fascicle shortening velocities in a preset range of cadences are not affected by changing crank power requirement, there was an increase in shortening velocities at the SSC as power output increased. This is in contrast with previous reports that shortening velocities were consistent with maximal power production at the SSC during low power output. Further, changes in joint power distributions were not observed in submaximal to maximal cycling transitions. Instead, cyclists selected higher cadences, which may reduce force requirements, but may also increase the internal work to coordinate the limbs and segmental accelerations for power production. At low power demands, our results suggest that minimization of muscle activation (EMG RMS) in the knee may be a component to consider when selecting a cadence. Thus, these findings highlight the important differences in mechanical and muscular activation responses to changes in power pedal demands during cycling, suggesting that muscle activation may be a critical aspect for riders in selecting their cadences, but the performance criterion that governs bicycling tasks remains unclear.

## Acknowledgments

The authors would like to acknowledge Vicente Riveros for his assistance with 3D figure design.

## Authors’ contributions

C.R-M., T.J.C, M.J.C. and G.A.L. conceptualized and designed the experiment. C.R-M. conducted the experiments. C.R-M analyzed data, prepared figures, and drafted manuscript. C.R-M., T.J.C., M.J.C. and G.A.L. interpreted results, revised manuscript, and approved final version of manuscript.

